# Periplasmic oxidized-protein repair during copper stress in *E. coli*: a focus on the metallochaperone CusF

**DOI:** 10.1101/2022.04.01.486667

**Authors:** Alexandra Vergnes, Camille Henry, Gaia Grassini, Laurent Loiseau, Sara El Hajj, Yann Denis, Anne Galinier, Didier Vertommen, Laurent Aussel, Benjamin Ezraty

**Author notes:** Department of Biochemistry, University of Wisconsin-Madison, USA.

## Abstract

Methionine residues are particularly sensitive to oxidation by reactive oxygen or chlorine species (ROS/RCS), leading to the appearance of methionine sulfoxide in proteins. This post-translational oxidation can be reversed by omnipresent protein repair pathways involving methionine sulfoxide reductases (Msr). In the periplasm of *Escherichia coli*, the enzymatic system MsrPQ, whose expression is triggered by the RCS, controls the redox status of methionine residues. Here we report that MsrPQ synthesis is also induced by copper stress via the CusSR two-component system, and that MsrPQ plays a role in copper homeostasis by maintaining the activity of the copper efflux pump, CusCFBA. Genetic and biochemical evidence suggest the metallochaperone CusF is the substrate of MsrPQ and our study reveal that CusF methionines are redox sensitive and can be restored by MsrPQ. Thus, the evolution of a CusSR-dependent synthesis of MsrPQ allows keeping copper homeostasis under aerobic conditions by maintenance of the reduced state of Met residues in copper-trafficking proteins.

**AUTHOR SUMMARY:** Our study investigates the interconnection between the copper stress response and the methionine redox homeostasis in the Gram-negative bacterium *Escherichia coli*. We report that the copper-activation of the CusSR two-component system induces the expression of the periplasmic oxidized-protein repair system, MsrPQ. We demonstrate that MsrPQ is crucial for CusCFBA copper efflux pump activity under aerobic conditions as it maintains the periplasmic component CusF in its functional reduced form. Methionine emerges as a critical residue in copper trafficking proteins: however this naturally-selected advantage must be balanced by methionine’s high susceptibility to oxidation. Therefore the induction of MsrPQ by copper allows copper homeostasis under aerobic conditions, illustrating that *E. coli* has developed an integrated and dynamic circuit for sensing and counteracting stress caused by copper and oxidants, thus allowing bacteria to adapt to host cellular defences.

## INTRODUCTION

Accumulation of damaged proteins hampers biological processes and can lead to cellular dysfunction and death. Chaperones, proteases and repair enzymes allow cells to confront these challenges and regulate protein homeostasis. The activity of these protein families defines “protein quality control” [1] and under stress conditions (high temperatures, oxidative or metal stress), signal transduction cascades up-regulate protein quality control to reduce the appearance of aggregation-prone molecules [2]. Protein quality control is also involved in housekeeping functions in different cellular compartments throughout the cellular life cycle. Within proteins, sulfur-containing amino acids such as methionine (Met) are targets for reactive oxygen species (ROS) and reactive chlorine species (RCS), the latter being more efficient at converting Met to its oxidized form, methionine sulfoxide (Met-O) [3]. This oxidation reaction is reversible due to the action of methionine sulfoxide reductases (Msr) [4]. MsrPQ in *E. coli* is an Msr system necessary for periplasmic proteins quality control, in which MsrP reduces Met-O and MsrQ is the membrane-bound partner required for MsrP activity [5]. We have shown that *msrPQ* expression is induced by RCS (HOCl) in a HprSR-dependent manner. HprSR is a two-component system (TCS), in which HprS is a histidine kinase (HK) sensor and HpsR the cytoplasmic response regulator (RR) [6]. The periplasmic chaperone SurA is one of the preferred substrates of MsrP [5] and proteomic studies have pinpointed processes including metal homeostasis, under the supervision of MsrP [5].

Copper is an essential prosthetic group in major *E. coli* enzymes, including cytochrome *bo* quinol oxidase and copper-zinc superoxide dismutase, however, high copper concentrations are toxic to the cell [7]. In aerobiosis, copper toxicity may be due to its involvement in the Fenton-like reactions which generate the highly reactive hydroxyl radicals (HO°) [8]. The Imlay group showed that the copper-mediated Fenton reaction does not cause oxidative DNA damage in *E. coli* cytoplasm [9]. Conversely, copper EPR spectroscopy suggested that most of the copper-mediated HO° formation does not occur near DNA, but in the periplasmic compartment [9]. Copper is more toxic under anaerobic conditions [10] and Fe-S clusters are the main intracellular targets of copper toxicity, even in the absence of oxygen [11]. Regulation of copper homeostasis is therefore required to maintain intracellular copper at low levels [12]. In *E. coli*, at least three systems are involved in copper tolerance: (i) CopA, a P-type ATPase which pumps copper from the cytoplasm to the periplasm [13]; (ii) CueO, a periplasmic multi-copper oxidase that oxidizes Cu(I) to the less toxic Cu(II) [14,15] and (iii) CusCFBA, an RND-type (<u>r</u>esistance, <u>n</u>odulation, <u>d</u>ivision) efflux pump responsible for extrusion of copper into the extracellular environment [16]. This RND-type efflux pump consists of CusA, the inner-membrane proton antiporter, CusB, the periplasmic protein, CusC the outer-membrane protein, and CusF, the periplasmic metallochaperone that supplies copper to the pump. The *cusCFBA* operon is under the control of the CusSR pathway in which CusS is the sensor and CusR the RR [17]. Finally the CueR transcriptional regulator regulates both *copA* and *cueO* expression [18].

Several lines of evidence point to the role of methionine residues in copper coordination within proteins such as CopA, CueO, CusF and CusAB proteins [7,19]. Mutation of the conserved Met204 in CopA yields an enzyme with a lower turnover rate, which is explained by a decrease in Cu(I) transfer efficiency from CopA to the chaperone CusF [20]. CueO has a methionine-rich helix which allows Cu(I) binding to provide a cuprous oxidase function [14,15,21]. The periplasmic copper chaperone CusF binds Cu(I) via two important methionine residues [16,22,23]. Also for the periplasmic adaptor CusB, and the inner membrane component CusA, methionine residues play a pivotal part in Cu(I) binding and in the stepwise shuttle mechanism by which the pump extrudes copper from the cell [24–27]. In summary, in many cases Met residues have been identified as crucial for copper resistance.

ROS/RCS could impair the detoxification function of CueO, CopA and CusCFBA through the oxidation of Met residues; MsrP would then be required to reduce Met-O to allow proteins to recover their copper homeostatic functions. This postulate is reinforced by a study showing that the CusSR system up-regulates the expression of the *hiuH* gene, located upstream of *msrP* [28,29], opening up the possibility that MsrP is produced during copper stress to maintain at least one of the three systems involved in copper tolerance. Here we report that *msrP* is induced during copper stress via CusSR, we then establish by a phenotypic approach that MsrP is crucial for maintaining CusCFBA pump activity under aerobic conditions. By focusing on the periplasmic proteins CusB and CusF, we demonstrate that the metallochaperone undergoes post-translational Met-O modification after H_2_O_2_ treatment, affecting its activity, which can be restored by MsrPQ.

## RESULTS

### *msrP* expression is induced by CuSO_4_ in a CusSR-dependent manner

We have recently shown that the *E. coli* genes *hiuH, msrP* and *msrQ* belong to the same operon [6] and previous studies have indicated that copper induces *hiuH* expression [28]. To corroborate these observations, we investigated copper’s role in the production of MsrP: to measure the effect of copper on the *hiuH-msrPQ* operon, quantitative reverse transcription polymerase chain reaction (qRT-PCR) experiments were performed in a wild-type strain of *E. coli* under aerobic conditions. Our results show that *hiuH*, *msrP* and *msrQ* mRNA levels increased significantly in copper-treated cells (~190, ~13 and ~30-fold respectively) (Fig. 1A). Western blot analyses showed higher MsrP protein levels following CuSO_4_ treatment (Fig. 1B). We then decided to investigate whether the TCS HprSR or CusSR regulates the expression of the *hiuH-msrPQ* operon under copper stress. The translational *msrP-lacZ* reporter fusion was introduced into the wild-type, Δ*hprRS* and Δ*cusRS* strains for ß-galactosidase assays. The strains carrying the chromosomal reporter fusion were cultured in M9/CASA medium in the absence or presence of 500 μM of CuSO_4_. The *msrP-lacZ* activity increased (≈ 4-fold) after exposure to CuSO_4_ in the wild-type and Δ*hprRS* strains, but not in the Δ*cusRS* strain. Our results demonstrate that the increase in *msrP* expression following copper exposure is dependent on CusSR but not on HprSR (Fig. 1C), consistently with previous reports [28]. The copper-dependent induction of *msrP* expression is lower than HOCl-HprSR dependent induction, in which the cells exhibited ≈ 60-fold higher ß-galactosidase activity (Fig. 1D and [5,6]). These results show that MsrP concentrations increase in response to copper in a CusSR-dependent manner.

**Figure 1.**
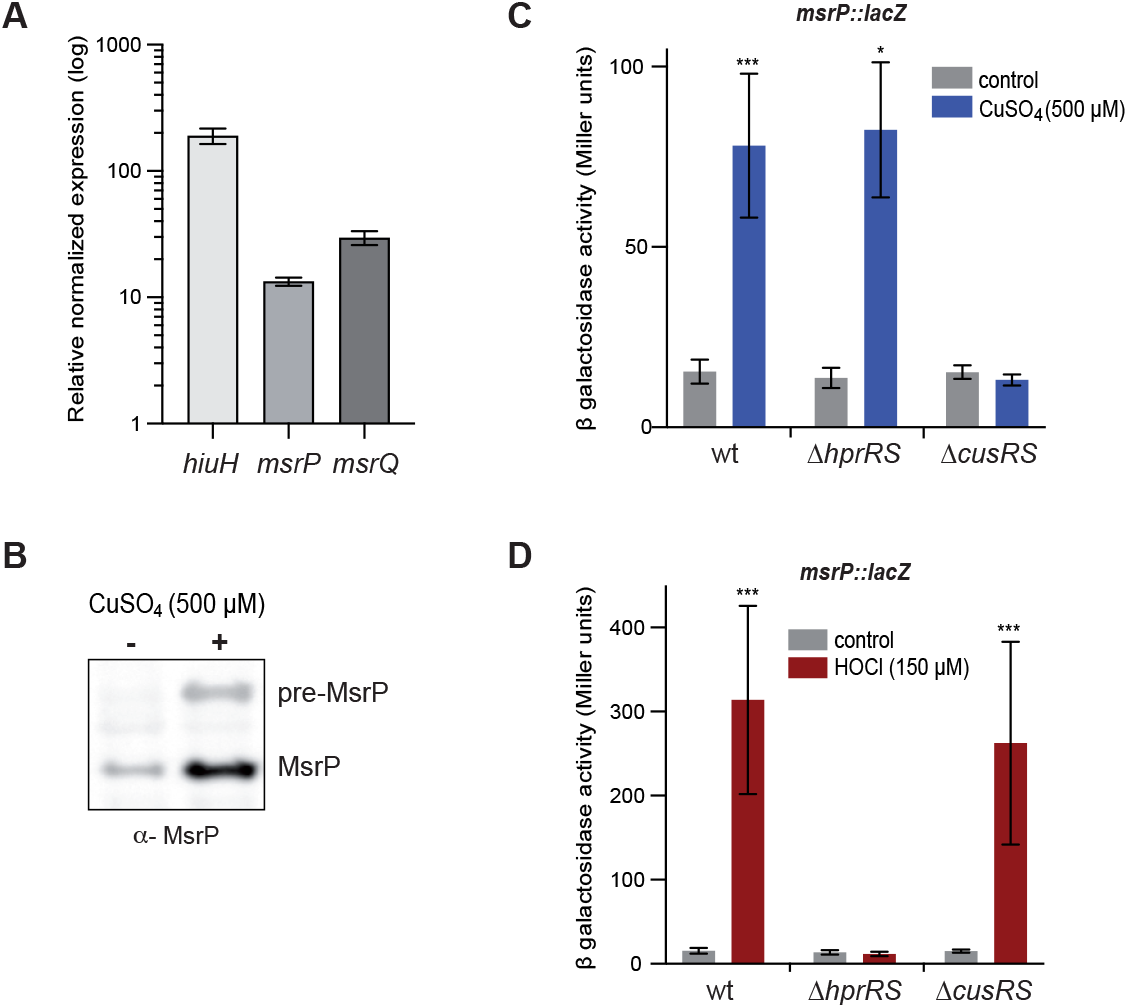
Copper regulates *msrP* expression in a CusSR-dependent manner. **A)** Relative normalized expression of *hiuH, msrP and msrQ* genes during copper stress. RNA was extracted from the wild-type strain grown in LB with CuSO_4_ (500 μM) to an OD_600nm_ ≈ 2. Quantitative real-time PCR was performed to amplify the *hiuH, msrP* and *msrQ* genes. Results are the means ± standard deviation of three independent experiments. **B)** Immunoblot analysis using an anti-MsrP antibody, showing the production of MsrP by CuSO_4_ (500 μM) stress. The image is representative of experiments carried out in triplicate. **C-D)** *msrP::lacZ* fusion was used as a proxy for *msrP* expression. Wild-type, Δ*hprRS* and Δ*cusRS* strains were grown in M9/CASA medium with or without the addition of 500 μM CuSO_4_ (**C**) or 150 μM HOCl (**D**), and ß-galactosidase assays were performed. Deletion of *cusRS* prevents *msrP* induction by copper, whereas deletion of *hprRS* prevents its induction by HOCl. Error bars, mean +/- s.e.m.; n=8 for wild-type and Δ*cusRS*, n=3 for Δ*hprRS*. Asterisks indicate a statistically significant difference between control and stressed conditions. *P ≤ 0.05; **P ≤ 0.01; and ***P ≤ 0.001 (Mann-Whitney U test).

### MsrP is required for copper tolerance

We hypothesized that MsrP might be important for cell growth when copper availability is high as *msrP* is part of the CusSR regulon. To test this, we exposed the Δ*msrP* strain to copper stress. The growth of the *msrP* mutant strain was first assayed on M9 plates containing CuSO_4_ (12.5 to 20 μM). Disruption of *msrP* did not lead to significant copper sensitivity compared to a wild type strain under aerobic growth conditions (Fig. 2A). We reasoned that functional redundancy between copper homeostasis systems might mask the importance of MsrP in copper tolerance (Fig. 2B). We therefore decided to focus on MsrP and the CusCFBA efflux pump, as they are both part of the CusSR-mediated response. To test our hypothesis, we introduced the Δ*copA* and the Δ*cueO* mutations into the Δ*msrP* mutant. The copper sensitivity of this triple mutant was monitored on M9 plates containing 5 μM of CuSO_4_. The Δ*copA* Δ*cueO* Δ*msrP* strain is more sensitive to copper than the parental Δ*copA* Δ*cueO*, MsrP proficient strain (Fig. 2C). We found that the copper sensitivity of the Δ*copA* Δ*cueO* Δ*msrP* strain was similar to that of the triple copper tolerance system mutant: Δ*copA* Δ*cueO* Δ*cusB* strain. These results suggest that MsrP may play a role in copper tolerance.

**Figure 2.**
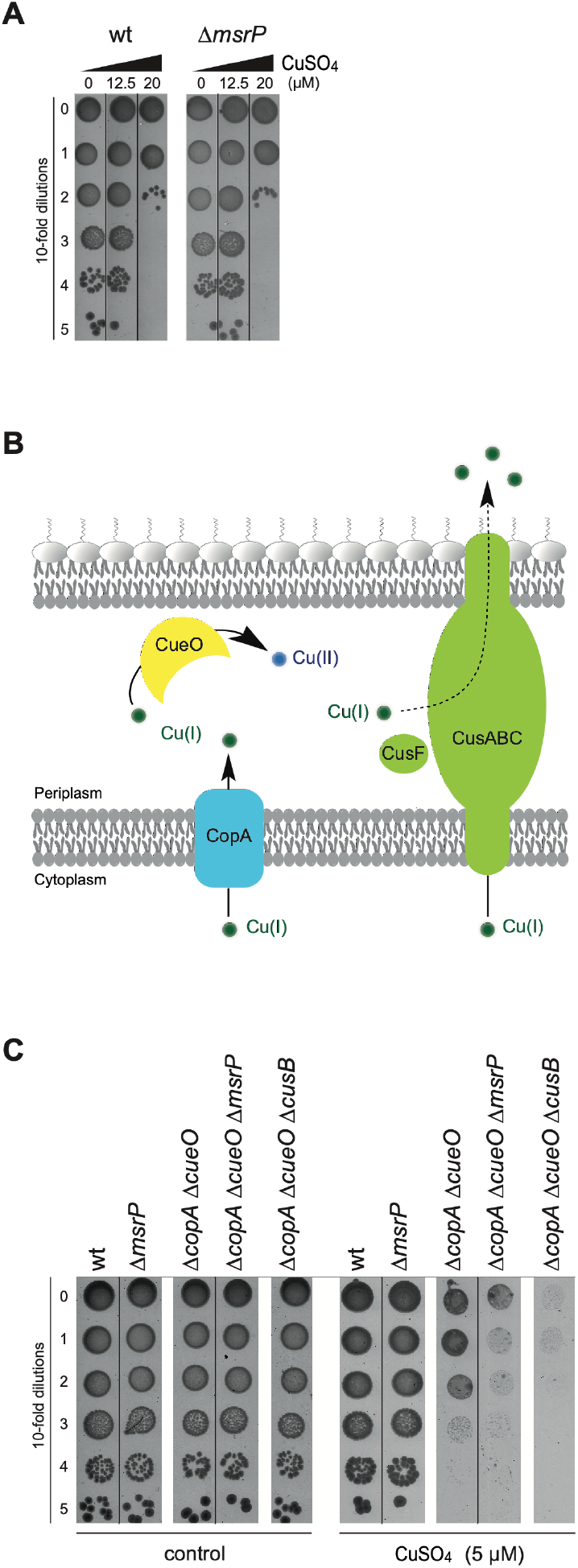
Involvement of MsrP in copper tolerance in *E. coli*. **A)** Plating efficiency of wild-type and Δ*msrP* strains in the presence of CuSO_4_. Cells were grown to an exponential phase (OD_600nm_ ≈ 0.1) at 37 °C in M9 medium and 10-fold serial dilutions were spotted onto M9 plates, with or without the addition of CuSO_4_ at the concentrations given (top panel). No significant difference was observed between either strains. **B)** Schematic view of the copper homeostasis systems in *E. coli*. CopA (in blue) translocates Cu(I) ions from the cytoplasm into the periplasm. CueO (in yellow) oxidizes Cu(I) ions to the less toxic form Cu(II). CusCBA efflux system (in green) pumps out copper to the extracellular environment. The CusF protein (in green), part of the *cusCFBA* operon, is a periplasmic metallochaperone which supplies copper to the pump. **C)** Plating efficiency of wild-type, Δ*msrP*, (Δ*copA* Δ*cueO*), (Δ*copA* Δ*cueO* Δ*msrP*) and (Δ*copA* Δ*cueO* Δ*cusB*) strains in the presence of CuSO_4_. Cells were grown to an exponential growth phase (OD_600nm_ ≈ 0.1) at 37 °C in M9 medium and 10-fold serial dilutions were spotted onto M9 plates, without stress (left side panel) and with CuSO_4_ (5 μM)(right side panel). The images are representative of experiments carried out at least three times.

### The copper sensitivity of Δ*copA* Δ*cueO* Δ*msrP* is oxygen-dependent

The above findings suggest that the periplasmic oxidized-protein repair system is part of the copper stress response. Thus, we investigated the possibility that the link between MsrP and copper was oxidative stress dependent by performing the copper sensitivity assay under anaerobic conditions. In doing so, we did not detect copper-dependent growth inhibition of the Δ*copA* Δ*cueO* Δ*msrP* mutant compared to the isogenic parental MsrP-proficient strain (Fig. 3A). To obtain more direct evidence that ROS are involved in the copper sensitivity of the strain lacking MsrP, we added an excess of catalase to plates before cell spreading - this method has been shown to reduce H_2_O_2_ levels under aerobic conditions [30]. Adding catalase to plates eliminated the *msrP*-deficient strain phenotype (Fig. 3B). These data are consistent with the copper sensitivity of the Δ*copA* Δ*cueO* Δ*msrP* strain being ROS dependent.

**Figure 3.**
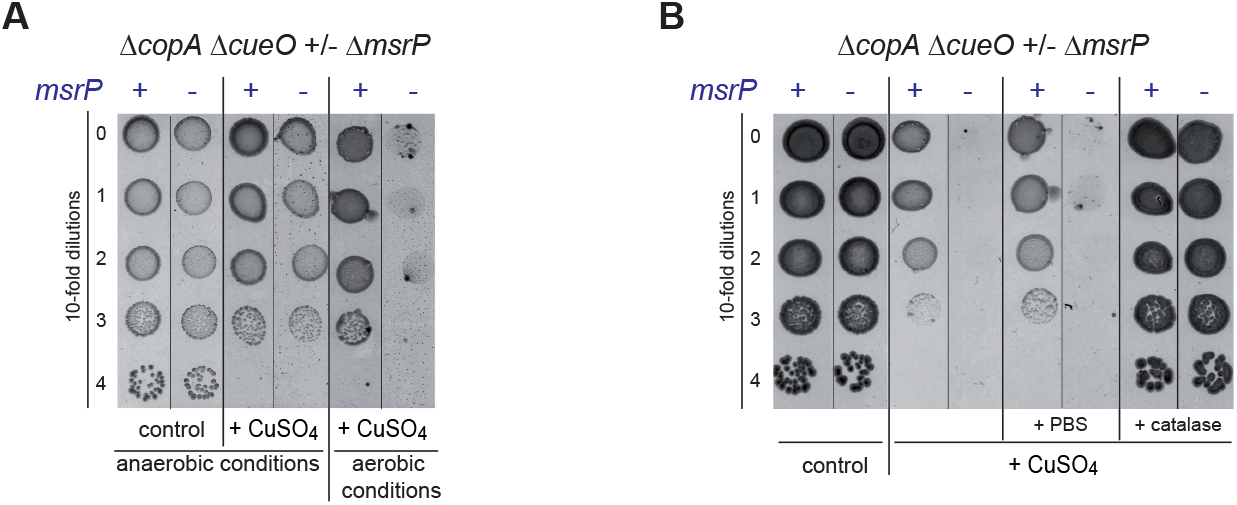
The copper hypersensitivity of the Δ*copA* Δ*cueO* Δ*msrP* strain is ROS dependent. **A)** Plating efficiency of Δ*copA* Δ*cueO* and Δ*copA* Δ*cueO* Δ*msrP* strains in the presence of CuSO_4_ (1.5 μM) under anaerobic conditions and in the presence of CuSO_4_ (5 μM) under aerobic conditions. The same protocol as described in Fig. 2 was used, except that plates were incubated in the absence of oxygen for 4 days. **B)** Plating efficiency of Δ*copA* Δ*cueO* and Δ*copA* Δ*cueO* Δ*msrP strains* in the presence of CuSO_4_ (25 μM) and catalase (2,000 units). 10-fold serial dilutions were spotted onto M9 plates with or without CuSO_4_ in the presence of PBS as a control or catalase under aerobic conditions. The images are representative of experiments carried out at least three times.

### MsrP is required for copper tolerance by maintaining CusF activity

Our above findings suggest that one or more components of the copper-efflux system CusCFBA may be damaged by oxidation. MsrP could therefore be essential for maintaining the CusCFBA pump in a reduced state. One prediction of our model is that CusCFBA pump overproduction should compensate for reduced efflux due to oxidation. To test this, we used the pCusCFBA plasmid encoding the whole operon and observed that the copper hypersensitivity of the Δ*copA* Δ*cueO* Δ*msrP* strain could be suppressed upon overexpression of the *cusCFBA* operon (Fig. 4) whereas the copper sensitivity phenotype of the Δ*copA* Δ*cueO* Δ*msrP* strain carrying the empty vector is less marked. In addition, we observed that the overexpression of *cusCFBA* genes is slightly harmful to the cell, even in the absence of copper (Fig. 4). To further test the prediction and to identify the limiting periplasmic component of the pump, we expressed the two periplasmic subunits CusB and CusF separately. We observed that overproduction of CusB, but not CusF, is toxic to the cell (Fig. 4). Interestingly, CusF overexpression in the Δ*copA* Δ*cueO* Δ*msrP* strain suppresses copper sensitivity of this strain (Fig. 4). In spite of our efforts to find a more discriminate assay, the difference between the mutant and the MsrP proficient strain were best observed on the agar-containing copper plate assay. Our results provide evidence that MsrP is at least involved in maintaining CusF activity.

**Figure 4.**
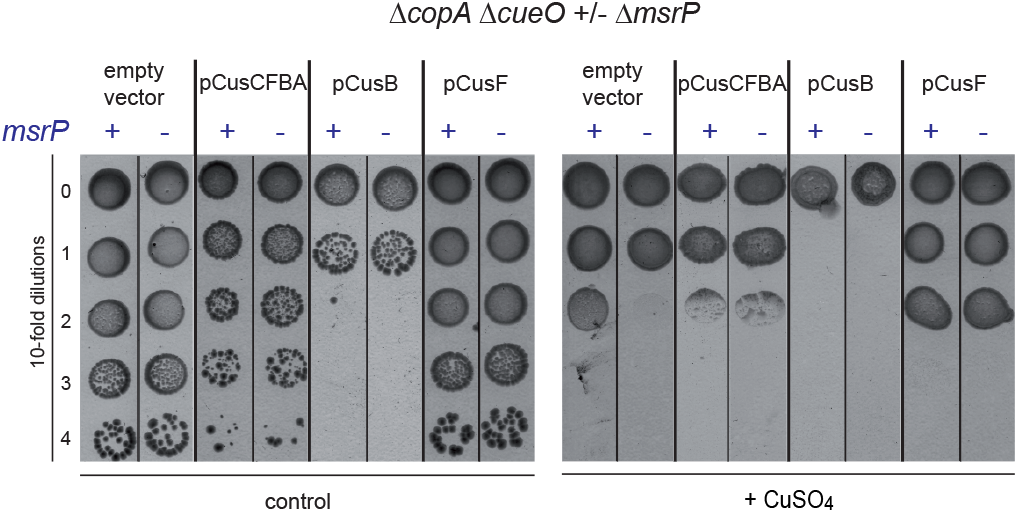
Overexpression of CusF suppressed the copper hypersensitivity of the Δ*copA* Δ*cueO* Δ*msrP* strain. Plating efficiency of the Δ*copA* Δ*cueO* Δ*msrP* strain carrying empty vector, pCusCFBA, pCusB or pCusF in the presence of CuSO_4_ (25 μM). The same protocol as described in Fig. 2 was used, except plates contained ampicillin (50 μg/ml) and IPTG (100 μM). The images are representative of experiments carried out at least three times.

### *In vivo* evidence for the consequences of CusF oxidation, using CusF^M47Q/M49Q^ as a proxy for of Met47 and Met49 oxidation

CusF is a soluble periplasmic protein that transfers copper directly to the CusCBA pump. The mature-CusF form contains four methionine residues (Met8, Met47, Met49 and Met59) of which Met47 and Met49, in addition to His36, are used as copper coordination ligands, with a nearby tryptophan (Trp44) capping the metal site [22]. Analysis of the apo-CusF structure shows that Met47 and Met49 are exposed to the solvent, with the accessible sulfur atoms accessible, whereas in its copper-bound form, CusF undergoes a conformational change whereby the sulfur of Met49 becomes inaccessible while Met47 appears to remain on the surface (Fig. 5A) [22,31]. We hypothesized that Met47 and Met49 oxidation could impair CusF activity and replaced these Met residues by Ile (I) or Gln (Q), the latter being a mimetic of Met-O [32]. We exploited the copper sensitivity of the Δ*copA* Δ*cueO* Δ*cusF* strain to assess the activity of the CusF variants by trans-complementation with the mutated genes (Fig. 5B). The phenotype of the Δ*copA* Δ*cueO* Δ*cusF* strain is complemented in trans by the gene expressing wild-type CusF. Conversely, expression of the CusF^M47I/M491^ does not complement the strain as previously reported [16]. The CusF^M47Q/M49Q^ variant partially complements the copper sensitivity of the Δ*cusF* strain but not as well as wild-type CusF (Fig. 5B) suggesting that Met47 and Met49 oxidation could hamper CusF activity.

**Figure 5.**
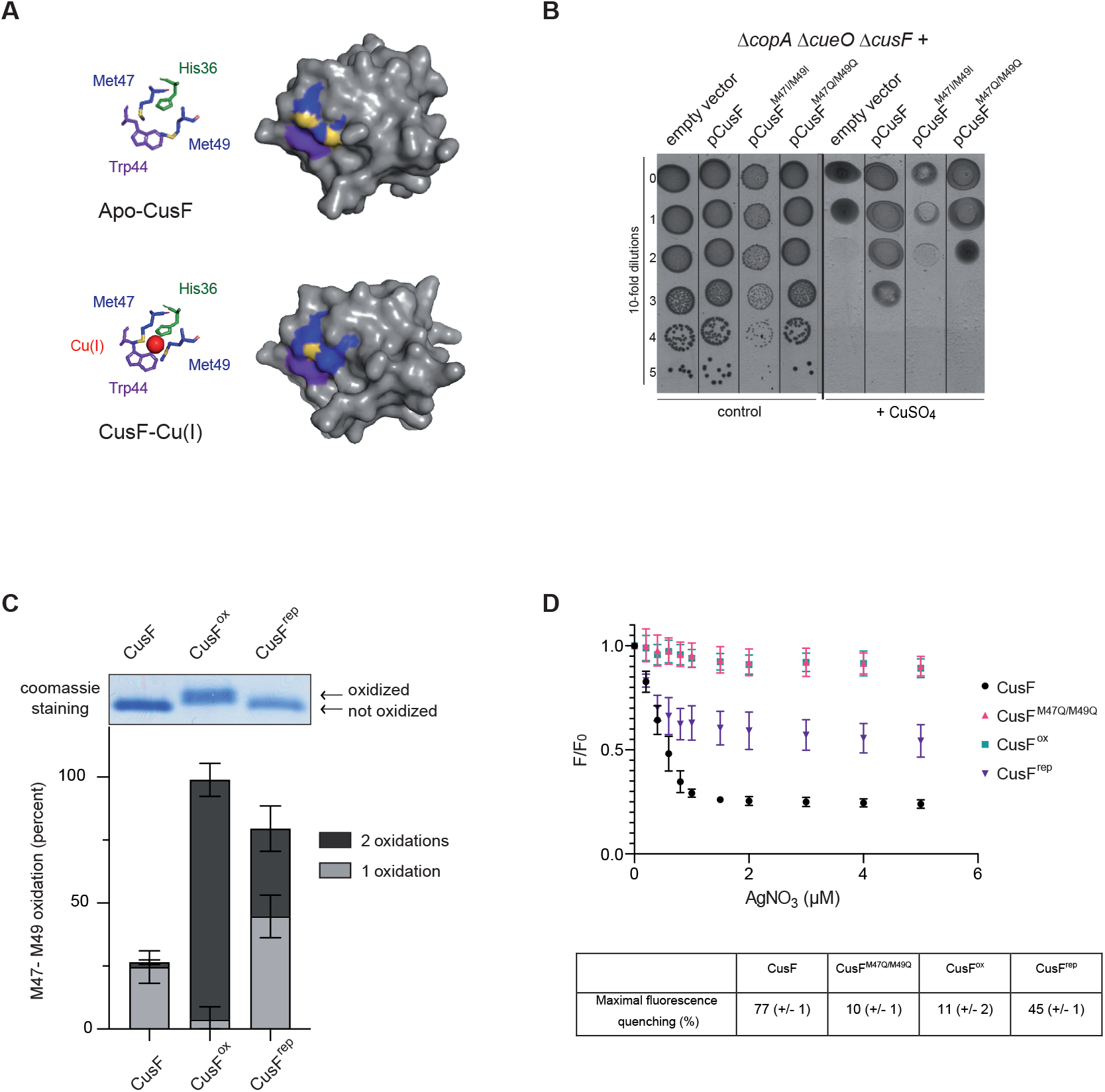
Methionine oxidation of CusF is deleterious. **A)** Aligned structures of the *E. coli* apo-CusF and CusF-Cu(I) adapted from PDB:1ZEQ and 2VB2 respectively [22,31] with stick and surface representations of CusF. Residues His 36 (green), Met 47, Met 49 (blue with sulphur atoms highlighted in yellow) and Trp44 (purple) are shown. The Cu(I) ion is shown in red. **B)** Plating efficiency of the Δ*copA* Δ*cueO* Δ*cusF* strain carrying empty vector, pCusF, pCusF^M47I/M49^ or pCusF^M47Q/M49Q^ vectors in the presence of CuSO_4_ (25 μM). The same protocol as described for Fig. 2 was used, except plates contained ampicillin (50μg/ml) and IPTG (50 μM). The images are representative of experiments carried out at least three times. **C)** Gel shift assay and mass spectrometry relative quantification by LFQ of the oxidation of Met47 and Met49. **D)** Silver binding analysed by quenching of intrinsic tryptophan fluorescence. Increasing concentrations of AgNO_3_ (0, 0.2, 0.4, 0.6, 0.8, 1, 1.5, 2, 3, 4, and 5 μM) were added to 1 μM CusF, CusF^M47Q/M49Q^, CusF^ox^ and CusF^rep^. The emission spectrum of CusF was recorded after each addition as described in the Materials and Methods. The integrated fluorescence peak (between 300 and 384 nm) in the presence of AgNO_3_ (F) was compared with the peak obtained in its absence (F_0_). The F/ F_0_ ratio was plotted against the concentration of AgNO_3_, after correction for the inner filter effect of AgNO_3_ measured on *N*-acetyltryptophanamide (NATA). The maximal fluorescence quenching for each variant of CusF was reported as a percentage in the table.

### Methionine oxidation of CusF gives rise to non-functional protein

We sought to characterize the metal-binding capacity of the CusF oxidized form. First, purified CusF protein was treated with H_2_O_2_ (50 mM) for 2 hours and analysed by mass spectrometry. CusF oxidation reaction can be first monitored by gel-shift assays (by SDS-polyacrylamide gel electrophoresis), as Met-O-containing proteins run slower than their reduced counterparts, leading to a mobility shift (Fig5C-upper panel) [33]. Met residues present in the mature CusF were identified in peptides detectable by mass spectrometry after trypsin digestion. Met47 and Met49 were part of the same peptide and therefore we could not determine the oxidation level of each residue separately. The Met47-Met49 containing peptide from untreated protein had around 25% of Met present as the Met-O form. This basal level of protein oxidation is commonly obtained and is usually assigned to the trypsin digestion protocol [34]. After H_2_O_2_ treatment, the proportion of Met-O increased to 98.9 % for Met47- and Met49-containing peptides (Fig. 5C-lower panel).

CusF has been shown to bind Cu(I) and Ag(I) with similar protein coordination chemistry [35,36]. We have taken advantage of the Ag(I)-binding property of CusF to assess the metal-binding capacities of the oxidized forms of CusF using AgNO_3_, instead of the highly toxic Cu(I) generation systems. For this we monitored the intrinsic fluorescence of CusF: Trp fluorescence emission peaks at 350 nm and the addition of increasing amounts of AgNO_3_ to CusF led to progressive fluorescence quenching (with a maximum of 77%). Compared to the CusF native form, the fluorescence quenching appeared different with the oxidized form of CusF (CusF^ox^): we observed a small decrease in intrinsic fluorescence even at the highest AgNO_3_ concentration tested (maximal fluorescence quenching = 11%). The difference could be due to the alteration of the Met residues involved in metal coordination ligands. To test this hypothesis, we purified the non-functional CusF^M47Q/M49Q^ variant, (substitutions mimicking Met47 and Met 49 oxidation), and measured its intrinsic fluorescence. Fluorescence quenching for CusF^M47Q/M49Q^ was comparable to that of CusF^ox^ (maximal fluorescence quenching of 10%), indicating that CusF oxidation disrupts the protein (Fig. 5D). We interpreted the low fluorescence quenching of CusF^ox^ and the mutated variant to be due to a Met-independent metal-binding domain. The fact that Trp residue emission is less affected by AgNO_3_ for both the oxidized and the mutated M to Q forms in comparison to the native protein could result from a local conformational change, leading to a non-functional protein. In order to test the reversibility of CusF oxidation, the CusF^ox^ was treated with MsrP enzyme in the presence of a reducing system (dithionite and benzyl viologen) to yield the repaired form (CusF^rep)^. Mass spectrometry and gel-shift assay of CusF^rep^ revealed a decrease in Met-O content, showing partial repair of CusF^ox^ (Fig. 5C). Fluorescence quenching was also partially restored (maximal fluorescence quenching of 45%, versus 77% for the native protein), probably reflecting a mix of oxidized and repaired CusF forms, in which the Trp residue has returned to its initial conformation (Fig. 5D). In conclusion, oxidized CusF is non-functional and MsrP can restore CusF activity by reducing Met-O.

## DISCUSSION

Methionine has emerged as a critical residue in copper trafficking proteins, providing labile sites that allow metal transfer. This selective advantage must be balanced with the high susceptibility of Met residues to oxidation. Indeed, under oxidative conditions (ROS, RCS), methionine is one of the preferred oxidation targets in proteins [37]. However, methionine oxidation is reversible due to the universal presence of the methionine sulfoxide reductases (MSR), which reduce oxidized methionine residues [3]. Here, we have demonstrated that in *E. coli*, the presence of copper induces the expression of the *msrP* gene encoding the enzyme involved in the repair of periplasmic oxidized proteins. Phenotypic analysis under aerobic conditions demonstrates the role of MsrP in maintaining the CusCFBA copper export pump. Genetic and biochemical analyses provide evidence that the oxidation of the CusF copper chaperone, at the very least, leads to the loss of function of this pump. In summary, (i) deletion of *msrP* is detrimental to CusCFBA activity, (ii) overexpression of *cusF* suppresses this phenotype, (iii) oxidized CusF contains Met-O residues, (iv) oxidized CusF is inactive as is mutated CusF M47Q/M49Q and (v) MsrP reduces Met in CusF and restores its activity

Interestingly, we show that both the *msrPQ* and *cusCFBA* operons are regulated by the CusSR TCS during copper stress: the existence of a common regulatory pathway for MsrP and CusF reinforces the idea of a functional link (Fig. 6). However, we cannot exclude the possibility that other components of the Cus pump are targeted by ROS/RCS, as well as the CopA and CueO proteins, which also contain methionine-rich sites involved in copper binding. In this study we used agar-plates containing copper under aerobic conditions -as copper defence systems depends on growing conditions [10], we can surmise that under other growth conditions MsrP maintains of CopA and/or CueO. Testing this hypothesis will be a field of future research.

**Figure 6.**
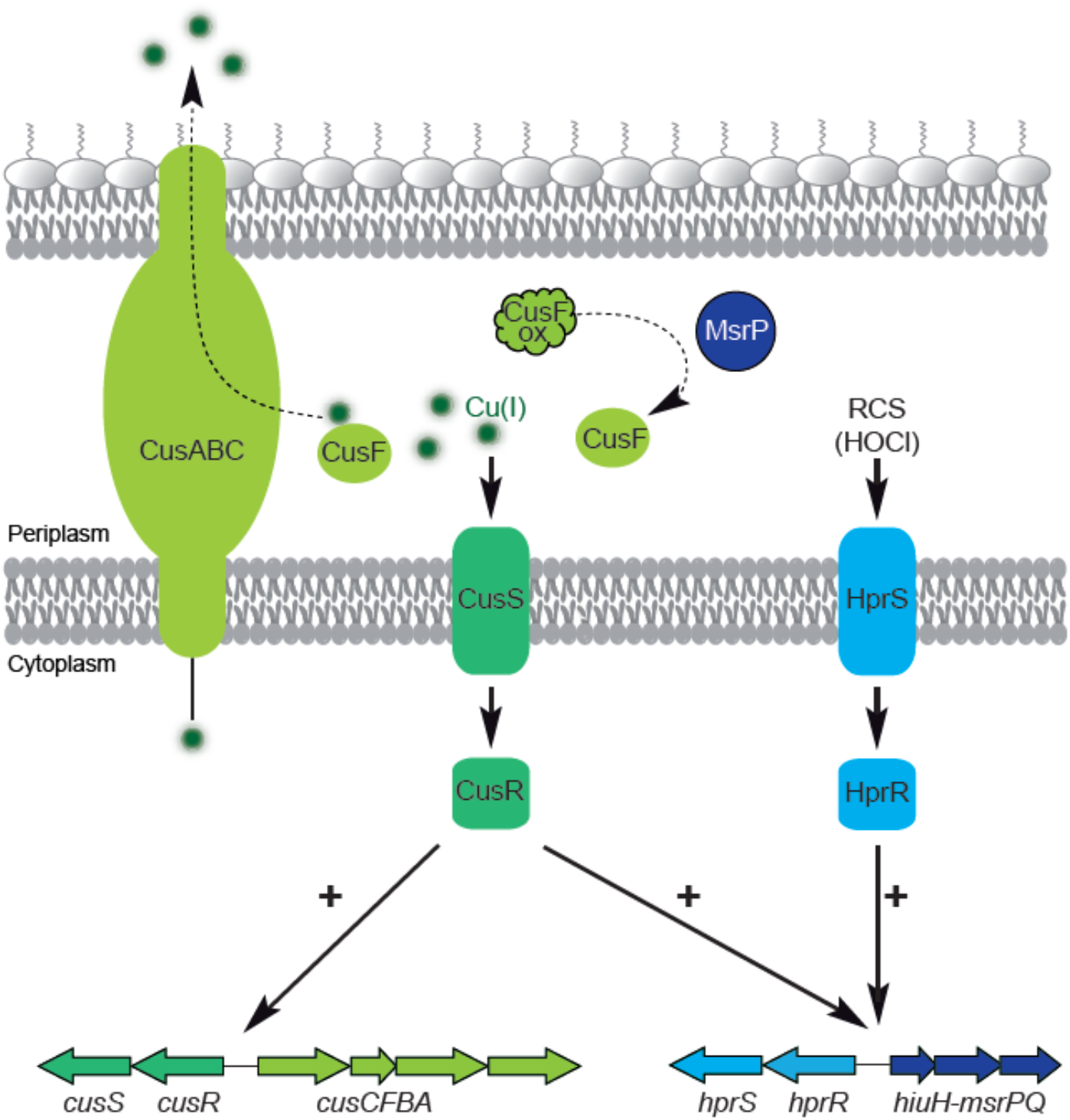
Copper efflux pump and oxidized-protein repair system are co-regulated. A working model illustrating the co-regulation of the copper efflux pump CusCFBA and the oxidized-protein repair system MsrPQ. Upon exposure to reactive chlorine species (RCS), the HprSR two-component system is activated leading to the up-regulation of the *hiuH-msrPQ* operon. Whereas, upon exposure to copper, the CusSR two-component system is activated leading to the up-regulation of the *cusCFAB* and *hiuH-msrPQ* operons. MsrP plays a role in copper homeostasis by controlling the redox status of methionine residues in the periplasmic metallochaperone CusF. CusF supplies copper to the efflux pump CusCBA, which then extrudes copper to the extracellular environment. By maintaining Met residues in a reduced form, MsrP appears to be essential for copper tolerance.

In this study, we show that in *E. coli* MsrP is expressed in the presence of copper. This observation could be explained by the fact that copper might participate in methionine oxidation via the copper-based Fenton reaction in the periplasm, like the analogous reaction driven by iron in the cytoplasm [9,38]. Our results reinforce this notion by demonstrating that even a protein involved in copper tolerance such as CusF is an oxidation target.

The *hiuH* gene, part of the *msrPQ* operon [6], encodes for a 5-hydroxyisourate (5-HIU) hydrolase, a protein involved in the purine catabolic pathway [39]. This enzyme catalyses the conversion of 5-HIU, a degradation product of uric acid, into 2-oxo-4-hydroxy-4-carboxy-5-ureidoimidazoline (OHCU). Based on the fact that copper has been shown to strongly inhibit the HiuH activity of *Salmonella* [40], Urano *et al*. proposed that the copper–dependent transcriptional regulation of *hiuH* might be important in maintaining uric acid metabolism [29]. Uric acid is generally considered to be an antioxidant having a free radical scavenging activity, but an opposite role as a copper-dependent pro-oxidant has also been reported [41]. Consequently, another hypothesis is that the copper and ROS/RCS up-regulation of *hiuH* might have a physiological role during oxidative stress. HprR and CusR have been shown to have the same recognition sequence and can bind to the consensus box with different affinities [29], leading to a collaborative or competitive interplay depending on the concentration of regulatory proteins. Therefore, a better characterization of the cross-regulation (copper *versus* oxidative stress) by the two TCS appears necessary. ROS/RCS and copper stresses are encountered during host-pathogen interactions [42]. During infection, phagocytic cells such as neutrophils produce ROS/RCS through NADPH oxidase and myeloperoxidase and accumulate copper in their phagosome via the ATP7A pump [43]. Thus, pathogens face both stresses at the same time. The interconnection between antimicrobial compounds produced by the immune system, like copper and HOCl, are an under-explored subject. Recently, the Gray laboratory reported that copper protects *E. coli* against killing by HOCl [44]. They identified the Cu(II) reductase RclA, which is induced by HOCl stress, as a central HOCl/copper combination resistance actor. The authors proposed that RclA prevent the formation of highly reactive Cu(III) by limiting the amount of Cu(II). Therefore, copper redox chemistry appears to be critical in the interaction between bacteria and the innate immune system. Interestingly, our study shows that *E. coli* has developed an integrated and dynamic circuit to sense and resist the combinatorial stresses caused by copper and HOCl, thus conferring an important adaptive capacity to host cellular defences. Our findings will be confirmed by future investigations examining the interplay between copper/HOCl stresses and protein oxidation during pathogenesis.

## ACKNOWLEDGEMENTS

We thank the members of the Ezraty group for comments on the manuscript, advice and discussions. Thanks to Pr. Dietrich H. Nies (Martin-Luther-Universitat Halle-Wittenberg) for providing CusF plasmids. We also thank M. Ilbert (BIP-CNRS), D. Byrne-Kodjabachian (IMM-CNRS) for helpful suggestions, reagents and comments on the manuscript. Special thanks to the former Marseillaise Barras team (Team Barras 4 ever) and to Frederic Barras (now at the Institut Pasteur) for lab space, support and discussions. A.V was supported by the Agence Nationale Recherche (ANR) (#ANR-16-CE11-0012-02 METOXIC), C.H. by the Fondation pour la Recherche Médicale (FRM) and G.G by AMidex (AMidex-Post-doc). This work was supported by grants from the ANR (#ANR-16-CE11-0012-02 METOXIC and #ANR-21-CE44-0024 MetCop) and CNRS (#PICS-PROTOX).

The authors declare no conflict of interest.

## MATERIALS AND METHODS

### Strains and microbial techniques

The strains used in this study are listed in Table 1. The corresponding alleles for of the deletion mutants were transferred from the Keio collection strain into the MG1655 wild-type strain by P1 transduction standard procedure and checked by PCR. The *hprRS* deletion mutant (strain CH100) was generated using a PCR knockout method developed by Datsenko and Wanner [45]. Briefly, a DNA fragment containing the *cat* gene flanked by the homologous sequences found upstream of the *hprR* gene and downstream of the *hprS* gene was PCR-amplified using pKD3 as template and the oligonucleotides *P1_Up_YedW*(*HprR*) and *P2_Down_YedV*(*HprS*). The fragment was transformed into strain MG1655, carrying plasmid pKD46, by electroporation. Chloramphenicol-resistant clones were selected and verified by PCR. The same procedure was used for the *cusRS* deletion mutant (strain GG100) with the oligonucleotides *cusS_kan_rev* and *cusR_kan_for*. Primer sequences used in this study are listed in Supplementary Table 2.

**Table 1.**
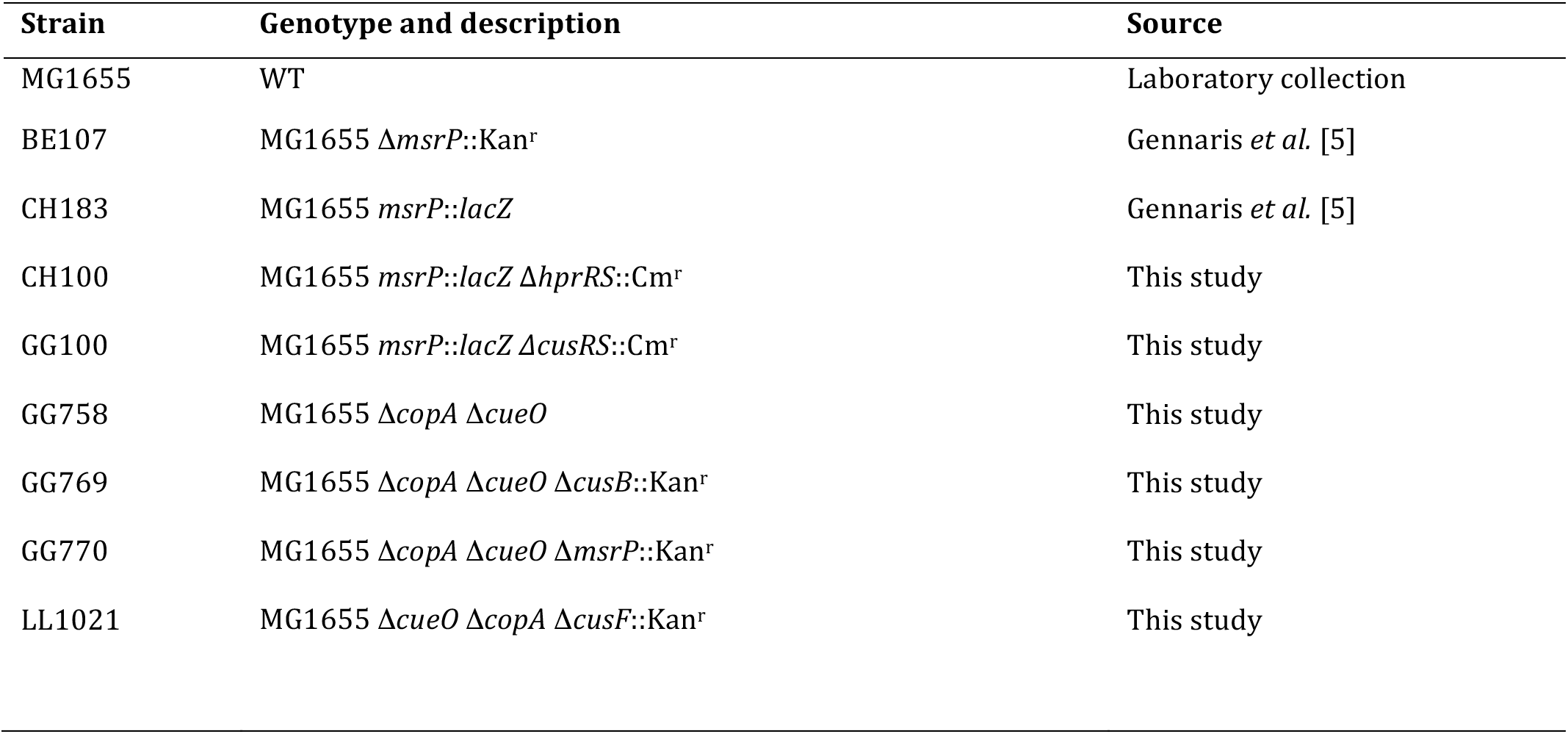
Strains used in this study. This table contains the information regarding the strains used in this study, including strain names, genotypes, description and source.

**Table 2.**
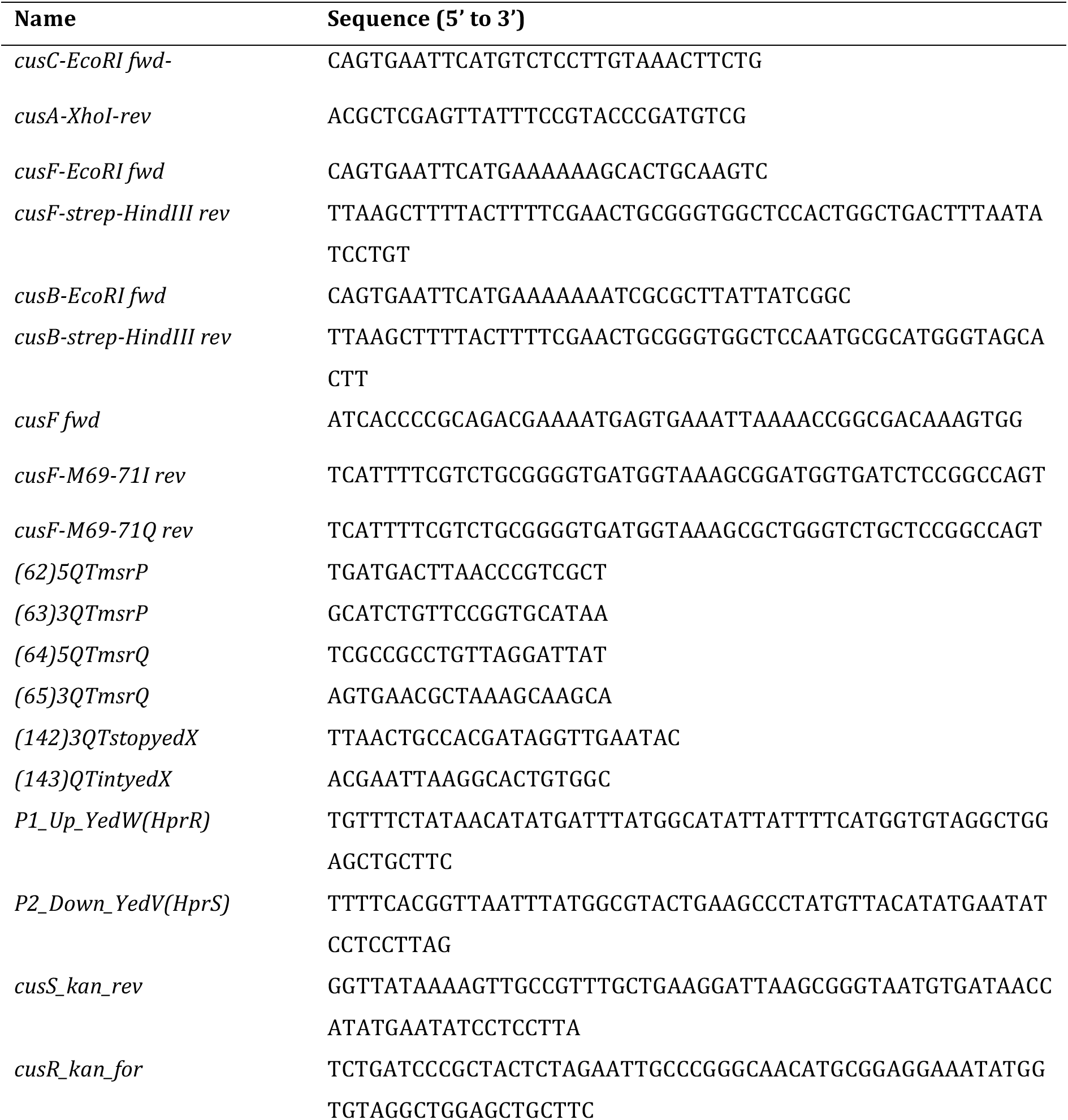
Primers used in this study. This table contains the information regarding the primers used in this study, including primer names and sequences.

### Plasmid construction

The plasmids used in this study are listed in Table 3. The CusCFBA (IPTG induced) expression vector was constructed by amplifying the *cusCFBA* operon was amplified from the chromosome (MG1655) using primers *cusC-EcoRI fwd* and *cusA-XhoI-rev*. The resulting PCR product was cloned into PJF119EH using *EcoRI* and *XhoI/SalI* restriction sites, generating plasmid pAV79.

**Table 3.** Plasmids used in this study. This table contains the information regarding the plasmids used in this study, including plasmid names, genotypes, description and source.

| Plasmid | Genotype and description | Source |
| --- | --- | --- |
| pJF119-EH | $P_{lac}$ promoter, IPTG inducible, Amp<sup>R</sup> selection | Laboratory collection |
| pAV79 (pCusCFBA) | pJF119-EH-CusCFBA | This study |
| pAV54 (pCusF) | pJF119-EH-CusF(Strep-TagII) | This study |
| pAV83 | pJF119-EH-CusF(Strep-TagII) M69I/M71I | This study |
| pAV84 | pJF119-EH-CusF(Strep-TagII) M69Q/M71Q | This study |
| pAV67 (pCusB) | pJF119-EH-CusB(Strep-TagII) | This study |
| pECD735 | pASK-IBA3plus CusF-StrepTagII | Dietrich H. Nies |
| pECD736 | pASK-IBA3plus CusF-StrepTagII M69I/M71I | Dietrich H. Nies |
| pAV96 | pASK-IBA3plus CusF-StrepTagII M69Q/M71Q | This study |

The CusF (IPTG induced) expression vector was constructed by amplifying the *cusF* gene from the chromosome (MG1655) using primers *cusF-EcoRI fwd* and *cusF-strep-HindIII rev*, which resulted in the fusion of a Strep-tag II coding sequence at the 3’ end. The PCR product was cloned into PJF119EH using *EcoRI* and *HindIII* restriction sites, generating plasmid pAV54. The CusB (IPTG-induced) expression vector was constructed using the same procedure, using primers *cusB-EcoRI fwd and cusB-strep-HindIII rev* and generating plasmid pAV67.

### *cusF* directed mutagenesis

50 μl PCR reactions were performed using Q5 Hot start High-Fidelity DNA polymerase (New England Biolabs), PJF119EH-cusF (pAV54) as the template and primers *cusF fwd* and *cusF-M69-71I rev* or *cusF-M69-71Q rev* (Supplementary Table 2). The resulting PCR products were digested using *DpnI*, purified using the GeneJET PCR purification kit (Thermo Fisher) and transformed into *E. coli* DH5α. Three colonies were randomly selected from each transformation, and the plasmids were isolated using the GeneJET Plasmid Miniprep kit (Thermo Fisher). DNA sequencing was carried out to assess the fidelity of the mutagenesis reaction.

### RNA preparation, PCR from cDNA and qRT-PCR

RNA from *E. coli* wild-type strain, grown to exponential growth phase (OD_600nm_≈ 2) at 37°C in LB supplemented or not with CuSO_4_ (500 μM), was extracted with Maxwell® 16 LEV miRNA Tissue Kit (Promega) according to the manufacturer’s instructions and was subjected to an extra TURBO DNase (Invitrogen) digestion step to eliminate the contaminating DNA. The RNA quality was assessed by a tape station system (Agilent). RNA was quantified at 260 nm using a NanoDrop 1000 spectrophotometer (Thermo Fisher Scientific). Quantitative real-time PCR analyses were performed on a CFX96 Real-Time System (Bio-Rad) in a final volume of 15 μl with 0.5 μM final concentration of each primer using the following program: 98 °C for 2 min, then 45 cycles of 98 °C for 5 s, 56 °C for 10 s, and 72 °C for 1 s. A final melting curve from 65 °C to 95 °C was added to determine amplification specificity. The amplification kinetics of each product were checked at the end of each cycle by measuring the fluorescence derived from the incorporation of EvaGreen into the double-stranded PCR products using the SsoFast EvaGreen Supermix 2X Kit (Bio-Rad, France). The results were analyzed using Bio-Rad CFX Maestro software, version 1.1 (Bio-Rad, France). RNA were quantified and normalized to the 16S rRNA housekeeping gene. qRT-PCR for each condition were carried out in triplicate. All biological repeats were selected and reported. Amplification efficiencies for each primer pairs were between 75 % and 100 %. Primer pairs used for qRT-PCR are listed in Supplementary Table 2.

### Immunoblot analysis of MsrP expression

To monitor MsrP expression levels after CuSO_4_ treatment, overnight cultures of wild-type cells (MG1655) were diluted to an OD_600nm_ of 0.04 in fresh LB medium (5 ml) and grown aerobically at 37 °C for 4 hours in the presence or absence of CuSO_4_ (500 μM). Samples were suspended in Laemmli SDS sample buffer (2 % SDS, 10 % glycerol, 60 mM Tris-HCl, pH 7.4, 0.01 % bromophenol blue), heated to 95 °C, and loaded onto an SDS-PAGE gel for immunoblot analysis. Protein amounts were standardized by taking into account the OD_600nm_ values of the cultures. Western blotting was performed using standard procedures, with primary antibodies directed against MsrP (rabbit sera; Jean-François Collet laboratory), followed by a horseradish peroxidase (HRP)-conjugated anti-rabbit IgG secondary antibody (Promega). Chemiluminescence signals were detected using the GE ImageQuant LAS4000 camera (GE Healthcare Life Sciences).

### Copper and HOCl induction assays

The *msrP::lacZ*-containing strains (CH183 (WT), CH1000 (Δ*hprRS*) and GG100 (Δ*cusRS*)) were grown at 37 °C under agitation in M9/CASA minimal medium. When cells reached an OD_600nm_ ≈ 0.2, cultures were split into three plastic tubes, one control tube, one containing 150 μM HOCl and one supplemented with 500 μM CuSO_4_, which were then incubated with an inclination of 90 ° with shaking at 37 °C. After 1 hour, 1 ml was harvested and the bacteria were resuspended in 1 ml of β-galactosidase buffer. Levels of β-galactosidase were measured as previously described [46].

### Copper survival assays

MG1655, BE107, GG758, GG769, GG770 and LL1021 cells were grown aerobically at 37 °C under agitation in 5 ml of M9 minimal medium in 50 ml conical polypropylene tubes (Sarstedt) with an inclination of 90°. When cultures reached OD_600nm_ ≈ 0.1, cells were harvested and diluted in PBS: 5 μL of 10-time serial dilutions were spotted onto M9 minimal medium-agar plates supplemented or not with CuSO_4_. Plates were incubated at 37 °C for 3 days. Ampicillin (50 μg/ml) and IPTG (100 μM) were added to solid and liquid media as required for plasmid selection. For the anaerobic conditions, the plates were incubated at 37 °C for 4 days in a BD GasPak system.

### Protein expression and purification

Wild-type CusF and variants were expressed and purified as previously described by Pr. Dietrich H. Nies laboratory [16]. MG1655 cells harboring plasmids pECD735, pECD736, pAV96 and over-expressing wild-type CusF, CusF^M69-711^ and CusF^M69-71Q^ proteins respectively, were grown aerobically at 37 °C in LB supplemented with ampicillin (200 μg/ml). When cells reached an OD_600nm_ of 0.8, expression was induced with anhydrotetracycline (200 μg/L final concentration) for 4 h at 30 °C. Periplasmic proteins were extracted and CusF was purified on a 5 ml StrepTrap HP column (GE healthcare) equilibrated with buffer A (10 mM NaPi, pH 8.0, 500 mM NaCl). After washing the column with buffer A, CusF was eluted with buffer A supplemented with desthiobiotin (2.5 mM). The fractions containing CusF were checked using SDS PAGE and the clean fractions were pooled and desalted with buffer 40 mM MOPS, pH7, 150 mM NaCl.

### Protein oxidation and repair *in vitro*

Wild-type CusF protein was oxidized using H_2_O_2_ (50 mM) for 2 hours at 37 °C. The reaction was stopped by buffer exchange using Zeba Spin Desalting Columns, 7K MWCO, with 40 mM MOPS, pH7, 150 mM NaCl. The CusF^ox^ protein formed was treated with MsrP enzyme in the presence of a reducing system to give the repaired form CusF^rep^ by incubating 100 μM CusF^ox^ for 2 hours at 30 °C in an anaerobic chamber with 4 μM purified MsrP, 10 mM benzyl-viologen and 10 mM dithionite. The reaction was stopped by buffer exchange using Zeba Spin Desalting Columns, 7K MWCO, with 40 mM MOPS, pH7, 150 mM NaCl.

### Mass spectrometry analysis

Samples were reduced and alkylated before digestion overnight with trypsin at 30 °C in 50 mM NH4HCO_3_ pH 8.0. Peptides were dissolved in solvent A (0.1 % TFA in 2 % ACN), immediately loaded onto a reverse-phase pre-column (Acclaim PepMap 100, Thermo Scientific) and eluted in backflush mode. Peptide separation was performed using a reverse-phase analytical column (Acclaim PepMap RSLC, 0.075 × 250 mm, Thermo Scientific) with a linear gradient of 4 %-36 % solvent B (0.1 % FA in 98 % ACN) for 36 min, 40 %-99 % solvent B for 10 min and holding at 99 % for the last 5 min at a constant flow rate of 300 nl/min on an EASY-nLC 1000 UPLC system. Peptide analysis was carried out using an Orbitrap Fusion Lumos tribrid mass spectrometer (ThermoFisher Scientific). The peptides were subjected to NSI source followed by tandem mass spectrometry (MS/MS) in Fusion Lumos coupled online to the UPLC. Intact peptides were detected in the Orbitrap at a resolution of 120,000. Peptides were selected for MS/MS using an HCD setting of 30; ion fragments were detected in the Orbitrap at a resolution of 30,000. The electrospray voltage applied was 2.1 kV. MS1 spectra were obtained with an AGC target of 4E5 ions and a maximum injection time of 50 ms, and targeted MS2 spectra were acquired with an AGC target of 2E5 ions and a maximum injection time of 60 ms. For MS scans, the m/z scan range was 350 to 1800. The resulting MS/MS data were processed and quantified by LFQ (area under the curve) using Proteome Discoverer 2.4 against an *E. coli* K12 protein database obtained from Uniprot. Mass error was set to 10 ppm for precursor ions and 0.05 Da for fragment ions. Oxidation (+15.99 Da) on Met, pyro-Glu formation from Gln and Glu at the peptide terminus, N-terminal removal of Met and acetylation were considered as variable modifications.

### Fluorescence measurements

All experiments were performed at 25°C using a SAFAS flx-Xenius 5117 spectrofluorimeter. Fluorescence measurements were carried out after dilution of wild-type CusF, CusF^ox^, CusF^rep^ or CusF^M69-71Q^ (1 μM final concentration) and equilibration for 5 min in 2 ml of a buffer containing 40 mM MOPS (pH=7) and 150 mM NaCl. Increasing concentrations of AgNO_3_ (0, 0.2, 0.4, 0.6, 0.8, 1, 1.5, 2, 3, 4, and 5 μM) were added and the emission fluorescence was scanned in the range of 300 to 384 nm, upon excitation at 284 nm. All spectra were corrected for buffer fluorescence with the same ligand concentration. Corrections for the inner-filter effect of the ligands were performed under the same conditions by using *N*-acetyltryptophanamide (NATA). The CusF fluorescence spectrum is centred at 350 nm and NATA spectrum at 357 nm. Peak integration was carried out for each ligand concentration.

### Statistical analysis

Mann-Whitney U tests were performed using the QI-Macros software (KnowWare International, Inc., Denver, CO).

